# Impact of thermal stress during pupal development in a key pollinator

**DOI:** 10.1101/2025.03.26.645459

**Authors:** Sandra Laußer, Christoph Kurze

## Abstract

The increasing frequency and intensity of extreme thermal events like heatwaves, driven by climate change, threaten global biodiversity and entire ecosystems. Despite alarming projections of population declines for key pollinators like bumblebees, little is known about the direct impact of varying elevated temperatures during their pupal development. To gain a better understanding of the dose-response to thermal stress, we experimentally exposed *Bombus terrestris* L. (Hymenoptera: Apidae) pupae *in vitro* to varying thermal stress lengths (3-7 days) and intensities (36-38°C). This mechanistic approach allowed us to uniquely isolate the direct physiological effects of temperature on pupal development from complex colony dynamics and assess individual emergence success, developmental duration, body size, relative lipid content, and wing geometrics morphometrics. At the critical threshold of 38°C, pupal emergence was drastically reduced (2-5 fold) and wing deformities increased (4-7 fold). Even excluding wing deformities, we found thermal stress-induced effects on emerged worker wing shape and directional asymmetry. Hence, our findings suggest that increased extreme thermal stress negatively impacts bumblebee pupal development. Given the relative heat tolerance of *B. terrestris*, we expect even greater vulnerability in more cold-adapted bumblebee species.

## 1. Introduction

Global warming is one of the most pressing challenges of our time [1, 2]. Particularly the increase in frequencies and intensities of heatwaves, defined as periods of abnormally hot temperatures sustained over at least three consecutive days, is concerning [3-6]. In fact, extreme thermal stress can severely impact a wide range of terrestrial animals such as insects, birds and mammals [4, 7, 8]. Despite recent advances in understanding the impact of thermal stress, our knowledge of the dose-response of extreme thermal stress (i.e. exposure variable temperatures and durations) on insect development and their physiological responses is still limited [7, 8].

Gaining a better understanding of the dose-response to thermal stress on insect physiology and fitness might be particularly important in threatened keystone species such as bumblebees (*Bombus* sp., Hymenoptera: Apidae). Bumblebees provide crucial pollination services in ecosystems and agriculture in the Northern hemisphere [9, 10]. As a cold-adapted taxon, bumblebees appear to be particularly vulnerable to climate change, which is reflected in their alarming population declines in some species [11-18]. However, heat tolerance varies significantly across species, with alpine and polar species being especially vulnerable to thermal stress [15, 19]. While ambient temperature fluctuations are buffered in ground nesting bumblebee species [20], those nesting above ground (e.g. *B. hypnorum* and *B. pascuorum* [15, 19]) and commercially managed species like *Bombus terrestris* L., widely used as crucial crop pollinators in greenhouses [21], face an increased risk of direct exposure to extreme thermal stress. Previous studies have shown that exposure to elevated temperatures impair cognition [22] and scent perception [23] in worker bumblebees. Under extreme thermal stress, the survival is significantly affected [24, 25]. Additionally, increasing temperatures negatively correlate with colony growth, which in turn negatively affects gyne production [26].

Colony fitness may not only be influenced by the quantity of offsprings [26], but also by their quality, expressed in their phenotypic traits. Wing morphology (shape, size and asymmetry) and storage reserves (e.g. fat body size) have been suggested as potential markers for monitoring bee health and pollination service, as they correlate with reproductive success, immunity, resilience and foraging efficiency [27]. While detailed dose-response data to environmental stressors at different life stages of bumblebees are still limited [28, 29], evidence suggest that offspring from colonies under constant thermal stress exhibit alterations in wing morphology, reduced body size and smaller antennae [30-33]. As a natural colony response to elevated nest temperatures worker spend significantly more time fanning to cool the brood than under optimal thermal conditions [34, 35]. As complex behaviroual responses can vary significantly between colonies, making it challenging to conduct detailed dose-response experiments under controlled conditions, *in vitro* experiments offer an alternative approach. These controlled experiments are particularly helpful to gain a better mechanistic understanding of the effects of environmental stressors at the precise developmental stages by uniquely isolating the direct physiological responses of individuals from the colony context at the cost of losing natural realism [28, 29].

While different developmental stages are likely differentially affected by thermal stress, it remains unclear which stages are more vulnerable and if short periods of thermal stress, like heatwaves, are sufficient to impair their development. To study the direct impact of thermal stress length and sevirity on pupal development in bumblebees, we used an *in vitro* rearing approach [28, 29]. This allowed us to precisely control the temperature by individual pupae, thus isolating their direct physiological and developmental responses from complex colony-level influences. We hypothesised that an increase in rearing temperatures and prolonged exposure would adversely affect the emergence rate, developmental duration, adult body size, fat body size, and wing development.

## 2. Methods

### (a) Experimental overview

To study the impact of thermal stress length and sevirity on pupal development in bumblebees, we collected 887 fourth instar larvae from 25 *Bombus terrestris* colonies to rear them *in vitro* following previous protocols [28, 29]. While our *in vitro* rearing temperature of 34°C [28, 29] exceeds typical nest temperatures [20], it aligns with brood temperatures, which are typically 2°C warmer than ambient nest temperatures [36, 37]. At the beginning of the pupal stage, pupae were pseudo-randomly assigned to one of the seven treatment groups (detailed in Table 1). We chose temperatures of 36°C, 37°C, and 38°C and durations of 3, 5, and 7 days to reflect potentially relevant extreme heatwave conditions observed in Europe [38-40]. Decision on thermal stress durations were based on our pilot study (unpublished data), which tested only 36°C for 3 days alongside controls and confirmed that an average pupation period would last about 8 days [29]. While we decided to expose pupae to 36°C for 5 and 7 days, we opted for shorter 3- and 5-day exposure durations for the treatments for 37°C and 38°C to avoid excessive pupal mortality at these more extreme temperatures and to focus on sub-lethal effects.

**Table 1.** Overview of the experimental design. This table details the rearing conditions for all experimental groups. Controls (Ctrl/34C) were reared in vitro at a constant temperature of 34°C until emergence as adults. Pupae of the thermal stress treatments experienced elevated temperatures (36°C, 37°C, and 38°C) for 3, 5 or 7 days at the start of pupation, followed by rearing at 34°C until emergence. Sample sizes (n) reflect the total number of pupae per treatment groups at the start of the experiment, but actual sample size for subsequent measurements were lower due to incomplete emergence.

| Treatment groups | Temperature | Duration | Sample size (n) |
| --- | --- | --- | --- |
| 34C/Ctrl | 34°C | until emergence | 127 |
| 36C5d | 36°C | 5 d | 79 |
| 36C7d | 36°C | 7 d | 80 |
| 37C3d | 37°C | 3 d | 75 |
| 37C5d | 37°C | 5 d | 73 |
| 38C3d | 38°C | 3 d | 64 |
| 38C5d | 38°C | 5 d | 106 |

We aimed to distribute pupae from the same collection date and source colony as broadly as possible among treatment groups (see Supplementary Table S1 for detailed information about collection cohorts, colony ID, and treatment assignment). Although a direct comparison between all treatment groups for each cohort was not feasible as we had access to only three incubators for four different rearing temperatures, our experimental design relied on a robust concurrent control group. These controls represent all collection cohorts and colonies as we continuously kept in one incubator at 34°C throughout the entire experiment. The other two incubators were rotated among 36°C, 37°C, and 38°C across experimental replications. This meant that not all treatment combinations/temperatures were run concurrently and did not derive from the same collection cohort (see Supplementary Table S1). To account for variability among colonies and collection cohorts, we included ‘colony ID’ and ‘collection date’ as a random effect into the statistical models whenever possible.

We checked their survival until adult bees emerged and measured their body mass. If adult bees emerged, we sampled them at age of 2 days and stored them at −20°C until subsequent morphometric measurements of their head width, intertegular distance (ITD), dry mass, and wings.

### (b) Colony husbandry

Larvae were collected from 25 queen-right colonies of *Bombus terrestris* (Natupol Research Hives, Koppert B.V., Netherlands) at various colony phases across multiple days between 19^th^ of June and 31^st^ of October 2023 to ensure a continuous supply and broad representation of colony cohorts (see Supplementary Table S1 for detailed collection dates, number of larvae collected per day, and their colony origin). Colonies were maintained under similar conditions to previous studies [28, 41]. In short, colonies were housed in the standard Natupol nest boxes (23 (l) x 25 (w) x 15 (h) cm), which were connected to foraging arenas (59 (l) x 39 (w) x 26 (h) cm) under a 14:10 h light:dark cycle. Bumblebees had *ad libitum* access to 70% (w/v) sucrose solution. Depending on colony size, each colony received 3-11 g of pollen candy daily, which consisted of 67% organic pollen (naturwaren-niederrhein GmbH, Germany), 25% sucrose (Südzucker AG, Germany), and 8% tap water. The room temperature was maintained at 25 °C ± 1 °C and 30-50% relative humidity.

### (c) Larvae collection and *in vitro* rearing

To collect 4^th^ instar larvae (L4), all adult bees from a colony were temporarily transferred to a separate cage. Then, L4 larvae were carefully removed from individual, spherical brood cells using soft tweezers. L4 larvae can be easily identified by the small opening for food provisioning (figure S1*a*). To minimize disturbance, colonies were only resampled after minimum period of at least two weeks.

Depending on the size of L4 larvae, they were carefully transferred into either 0.6 mL or 0.96 mL large 3D-printed polylactide (PLA) artificial brood cells (diameter of either 8 mm or 10 mm respectively, figure S1*b*). Those were then placed in 24- or 48-well clear flat bottom plates (Falcon/Corning, USA) (figure S1*c*) and kept inside a ventilated plastic container (18.5 x 18.5 x 11.5 cm) at 34°C (KB115, BINDER GmbH, Germany). A 120 mL cup of saturated sodium chloride solution was included within each container to maintain a relative humidity (RH) of 65 ± 10 %. The use of this setup facilitated the measurement of larval and pupal weight gain without the need for repeated handling them directly and allowed for the easy removal of any dead individuals.

L4 larvae were fed a pollen medium twice daily (morning and evening) until pupation and the start of the experiment. The pollen medium consisted of 50% w/v sucrose solution (Südzucker AG, Germany), 40% honeybee-collected organic pollen (Bio-Blütenpollen, naturwaren-niederrhein GmbH, Germany), 10% Bacto yeast extract (Bacto™, BD, USA), and 1% casein sodium salt from bovine milk (Sigma-Aldrich, Germany) [29]. The pollen medium was aliquoted and stored at −20°C until use. Before hand-feeding all L4 larvae, aliquots were warmed to 34°C and vortexed. For each 20-minute feeding session, well-plates containing the L4 larvae were placed on a heated plate at 35°C (Medax model 12801, Medax Nagel GmbH, Germany). Each larva initially received about 6 μL droplet (7.1 ± 1.6 mg, 10 μL micropipette tips had to be cut off at the 1 μL mark) of pollen medium applied to its ventral abdomen (figure S1*d*). Larvae that consumed all initial food received a second droplet. Satiation was determined when a larva curled up and stopped movement, while active larvae were considered hungry (Supplementary video 1). At the end of each feeding session, any remaining food was carefully removed to prevent tracheal blockage. Larvae were no longer fed when entering the pupal stage.

At the start of pupal stage, individuals were weighed (d = 0.1 mg, Sartorius AC120S, Sartorius AG, Germany) and pseudo-randomly assigned to one of seven treatment groups (detailed in Table 1), ensuring a balanced distribution across treatments and natal colonies (Table S1). Pupae were transferred to new humidity-controlled containers (65 ± 10 % RH), each containing a cup of saturated sodium chloride solution, inside an incubator set to the designated temperature. After the treatment period, pupae were returned to the control incubator at 34°C. We recorded the proportion of larvae that successfully reached adulthood.

### (d) Emerged adult bees

For each bee reaching adulthood, the emergence date was recorded to calculate the developmental time required to complete metamorphosis (i.e. pupal phase). Their body weight was directly measured using a precision scale (d = 0.1 mg, Sartorius AC120S, Sartorius AG, Germany). Adult bees were then housed individually in 25 mL snap-cap vial containing a cardboard bottom (approximately 5.5 x 1.5 cm) and mesh lid. Bees had *ad libitum* access to 60% (w/v) sugar solution via simple 1.5 mL feeder and kept at 25 °C ± 1 °C and 30-50% RH. Two-day-old adults, with fully developed and hardened wings, were frozen and stored at −20°C until further analysis.

### (e) Body size and lipid measurements

As a proxy for adult body size, the intertegular distance (ITD) and head width were measured using a digital microscope (VHX-500F, Keyence GmbH, Germany) [41]. In addition, their sex was determined by counting flagellomeres (workers have 10, males 11) [42]. We determined adult dry mass and lipid content following previous protocols [28, 41]. First, each specimen was carefully dissected by one cut at ventral abdominal segments from the stinger to the fourth sternite, then dried at 60°C for three days (U40, Memmert GmbH & Co. KG, Germany), and weighed (d = 0.1 mg, analytic balance M-Pact AX224, Sartorius GmbH, Germany). Subsequently, the body lipids were extracted using petroleum ether for five days. The ether was discarded, and the specimen were rinsed with fresh ether before being dried for an additional three days. The lipid content was calculated as the difference between the initial and post-extraction dry masses. The relative lipid content, serving as a proxy for the fat body size, was calculated as the percentage of lipid contents relative to the initial dry masses.

### (e) Wings

Wings were first detached at the tegulae and embedded side-by-side on a microscope slide (Roti^®^Mount, Carl Roth GmbH + Co. KG, Germany). Then, wing pairs were photographed at 20x magnification (Keyence VHX-500F). Two-dimensional coordinates of twenty landmarks [43] for each wing (figure S1*e*) were obtained using TpsDig2 (v2.32) [44]. While individuals with visibly undeveloped or fully deformed wings (figure S1*f*) were excluded from geometric morphometric analysis, we compared the proportion of individuals with developed, undeformed wings among treatment groups.

### (g) Statistical analyses

All statistical analyses and data visualizations were performed using R Statistical Software in R version 4.4.2 [45]. The complete code as RMD (R Markdown) output and datasets are provided in Dryad repository: https://doi.org/10.5061/dryad.n02v6wx7z.

We followed a hypothesis driven statistical approach. Whenever possible, we used generalized linear mixed effect models (GLMMs) using the *glmmTMB* package [46] to analyse effects of elevated temperatures and exposure duration, and their interactions, as fixed factors on emergence rates (i.e. pupae reaching adulthood), developmental time (i.e. days to emergence), adult mass, head width, ITD, relative lipid content, and wing development (i.e. either fully developed or deformed) with binomial (logit link), beta regression (logit link) or gamma (log link) data structures (see Supplementary RMD output for detailed model structure and selection procedure). We included ‘colony ID’ and ‘collection cohort’ (see Table S1) as random effects to account for colony and temporal-specific variability. Due to limited sample sizes for male specific data and unbalanced sample size distribution, we excluded male data and analysed the effects of thermal stress on adult mass, dry mass, ITD, and head width for worker data separately (the entire data are presented in figure S3). This approach was chosen to avoid sex-specific biases that could potentially mask treatment effects. In fact, we found sex-specific effects on adult mass (Kruskal-Wallis: *χ*^2^ = 99.723, df = 1, p < 0.0001), adult head width (*χ*^2^ = 88.556, df = 1, p < 0.0001), ITD (*χ*^2^ = 59.588, df = 1, p < 0.0001), and adult relative lipid content (*χ*^2^ = 6.748, df = 1, p < 0.01) (see Supplementary RMD output for a detailed analysis of the entire data).

For each response variable, a candidate set of GLMMs including potential explanatory variables was constructed to identify the most parsimonious and best-fitting model, based on the Akaike information criterion (AIC), with lower AIC values indicating a better fit (see Table S2 of a simplified overview selection procedure for models meeting model assumptions). The best models were compared with their respective null-models, including only random effects, using likelihood ratio tests (LRTs) to determine statistical significance of added terms. Model assumptions and dispersion of the data were checked using the *DHARMa* package [47]. Significance levels (p < 0.05) were determined using the *Anova* function when model assumptions were met. Due to our experimental design limitations described above, we specifically tested the effects of each discrete treatment combination compared to the controls using the function *emmeans* [48], adjusted with Holm-Bonferroni correction for multiple comparisons. When model assumptions of GLMMs were not met, treatment effects were tested with a non-parametric Kruskal-Wallis rank sum test, followed by post-hoc comparisons between each treated group with the controls using Dunn’s test with Holm-Bonferroni correction.

Geometric morphometric analyses of wing size, shape, and asymmetry on fully developed left and right forewings (i.e. excluding all deformed wings) of workers on the obtained TPS data using geomorph package [49], following a similar approach as previous studies [31, 43]. Due to sample size limitation of males, we also excluded male data from the geometric morphometric analyses of wings.

First, right wings were first mirrored to ensure landmark correspondence. Missing landmark data, representing morphological variation, were accounted for using the function *estimate*.*missing*. To remove non-shape variation, a Generalized Procrustes Analysis (GPA) was performed using *gpagen*. A Principal Component Analysis (PCA) was conducted on the Procrustes coordinates using *gm*.*prcomp*, PCA scores (PC1 and PC2) were calculated, and data distribution was visually assessed. Wings with PC1 z-scores exceeding ± 2 standard deviations were removed as outliers, where PC1 explained 78% of the data’s variation. Treatment groups with fewer than six individuals were also excluded from further analysis, which was only the case for workers from the 5-day-exposure at 38°C (38C5d).

Then, centroid size and mean centroid size were calculated for each wing and each bee, respectively, as a measure of wing size. To analyse shape variation, a permutational multivariate analysis of variance (PERMANOVA) was performed using the *adonis2* function. Homoscedasticity was evaluated using *betadisper*, and pairwise comparisons were conducted using *pairwise*.*adonis* with Holm-Bonferroni correction. Directional asymmetry (DA) was calculated as the difference between left and right wing Procrustes coordinates, and DA magnitude was calculated as the mean Euclidean distance between corresponding landmarks. To assess fluctuating asymmetry (FA), a bilateral symmetry analysis was performed using *bilat*.*symmetry* and calculating the FA magnitude as the Euclidean norm of the FA component. The effects of treatment on centroid size, DA and FA magnitude were tested using linear regression models (LM) when assumptions were met, followed by contrast pairwise comparisons for significant results as described above. Kruskal-Wallis tests were used as a non-parametric alternative, followed by Dunn’s post-hoc test to compare treatment groups to the control by applying Holm-Bonferroni corrections.

## Results

We found a strong interaction effect between thermal stress temperature and duration on the proportion of pupae reaching adulthood (GLMM: *χ*^2^ = 25.813, df = 1, p < 0.0001; figure 1*a*). This effect was mainly driven by increasing temperatures (*χ*^2^ = 24.549, df = 1, p < 0.0001). Pairwise comparisons revealed that pupae exposed to 38°C for 3 days were 43% (38C3d: z = 3.655, p < 0.01) less likely to reach adulthood compared to the controls. When exposed to 38°C for 5 days the emergence rate dropped by 83 % (38C5d: z = 7.236, p < 0.0001) compared to the controls.

**Figure 1.**
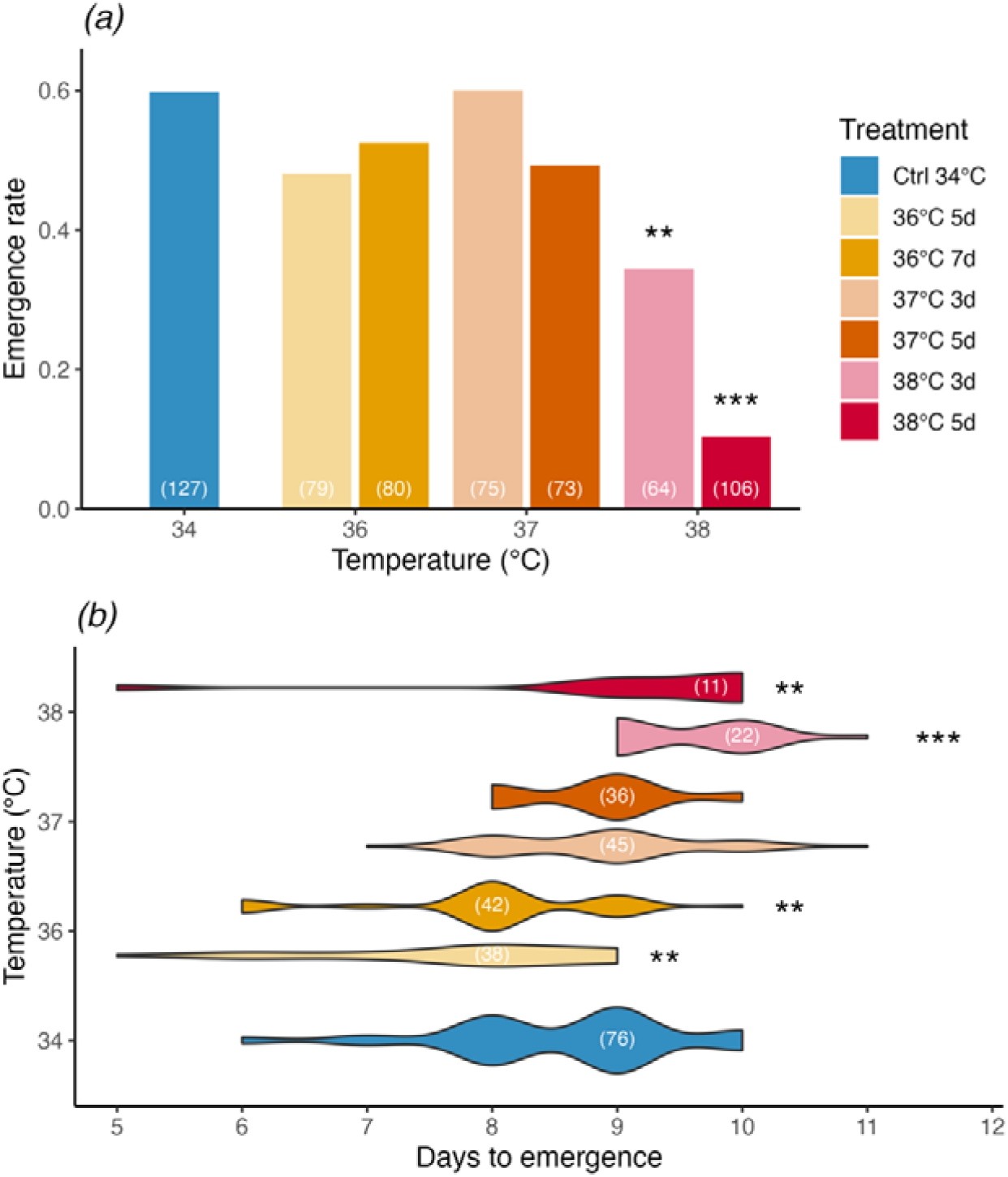
Direct effects of varying intensity of thermal stress on *(a)* emergence rate and *(b)* developmental duration in *B. terrestris* pupae. Controls (Ctrl, reared at constant 34°C, blue) were compared to individuals exposed to increased temperatures: of 36°C for 5 days (light warm yellow) and 7 days (warm yellow), 37°C for 3 days (light reddish-orange) and 5 days (reddish orange), and 38°C for 3 days (light red) and 5 days (deep red). Sample sizes are indicated by white numbers and significant differences are marked by asterisks (* p < 0.05, ** p < 0.01, *** p < 0.001).

Thermal stress intensity also significantly affected pupal developmental duration (Kruskal-Wallis: *χ*^2^ = 69.584, df = 6, p < 0.0001; figure 1*b*; worker and male data are presented separately in figure S2). While an exposure to 36°C shortened development compared to controls (Dunn’s test: 36C5d, z = 3.253, p < 0.01; 36C7d, z = 3.142, p < 0.01), an exposure to 38°C prolonged pupal development (38C3d, z = −4.393, p < 0.0001; 38C3d, z = −2.739, p < 0.01). There was no significant effect found for the 37°C exposure compared to controls (37C3d, z = −1.361, p > 0.5; 37C5d, z = −1.164, p > 0.5).

We found no significant effects of thermal stress treatments on worker body size proxies, including adult mass (Kruskal-Wallis: *χ*^2^ = 9.125, df = 6, p > 0.05; figure 2*a*) and ITD (Kruskal-Wallis: *χ*^2^ = 9.571, df = 6, p > 0.05; figure 2*c*>). While there was a weak significant effect on worker head width (Kruskal-Wallis: *χ*^2^ = 16.131, df = 6, p < 0.05, figure 2*b*), we found no significant differences between treatment groups and controls (Dunn’s tests: p > 0.05). However, female head width varied with temperature (Kruskal-Wallis: *χ*^2^ = 12.311, df = 3, p < 0.01), where workers exposed to 38°C as pupae had slightly wider heads than the controls (z = −2.227, p < 0.05).

**Figure 2.**
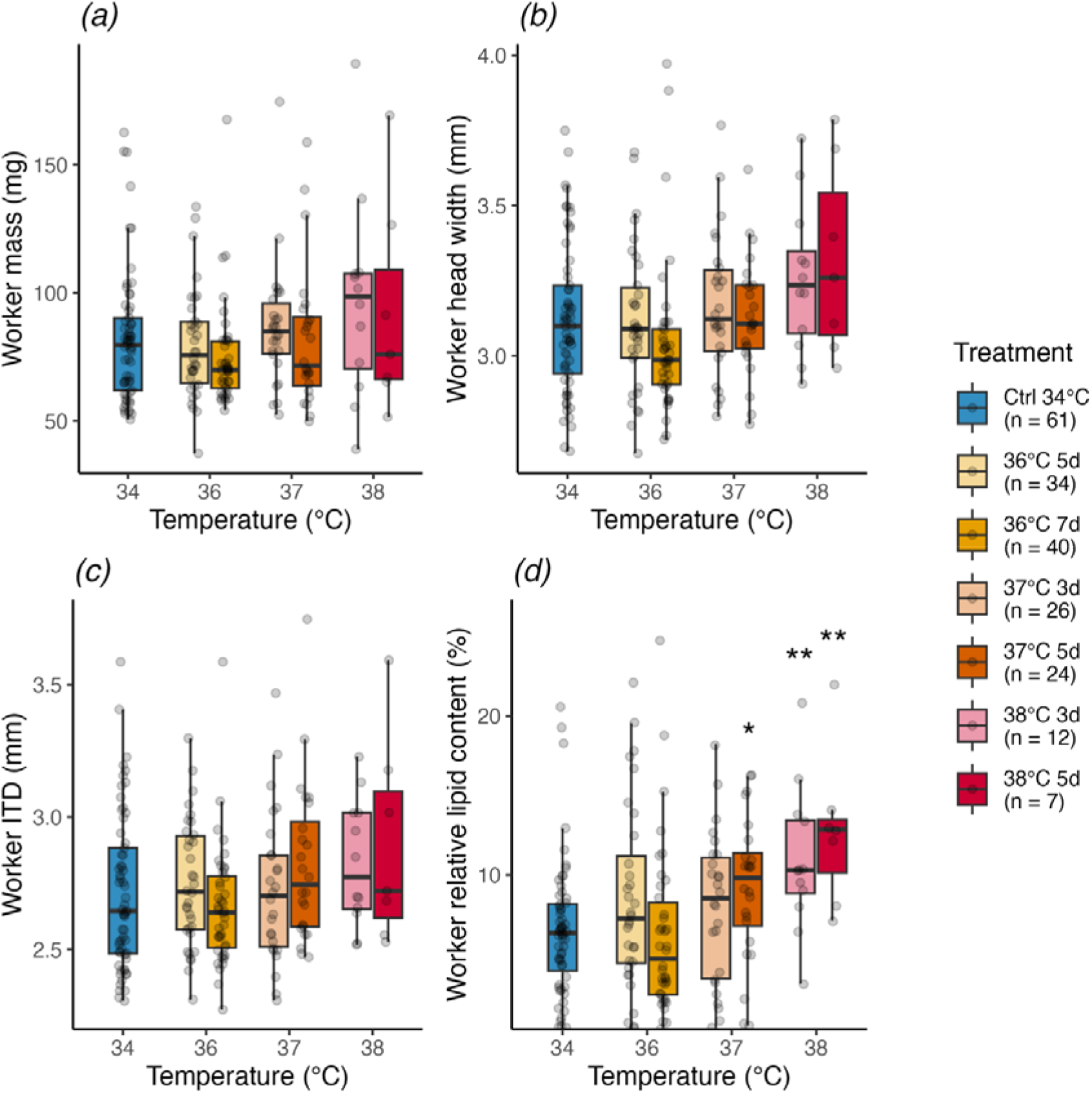
Effects thermal stress exposure during pupal development on adult worker morphometrics and lipid content in *B. terrestris*: *(a)* adult mass, *(b)* head width, *(c)* intertegular distance (ITD), *(d)* and relative lipid content (2-day-old adults). Controls (34°C, blue) were compared to individuals exposed to increased temperatures: of 36°C for 5 days (light warm yellow) and 7 days (warm yellow), 37°C for 3 days (light reddish-orange) and 5 days (reddish orange), and 38°C for 3 days (light red) and 5 days (deep red). Sample sizes (*n*) are provided in the legend. Asterisks denote significant differences (* p < 0.05, ** p < 0.01).

In contrast, we found that thermal stress temperature significantly influenced relative lipid content in adult workers (Kruskal-Wallis: *χ*^2^ = 25.847, df = 6, p < 0.001, figure 2*d*). Pairwise comparisons indicated significant differences between 37C5d (Dunn’s test: z = −2.592, p < 0.05), 38C3d (z = −2.844, p < 0.05), and 38C5d (z = −3.012, p < 0.01) and the controls (figure 2*d*).

We found that thermal stress temperature impacted wing development (GLMM: *χ*^2^ = 9.410, df = 1, p < 0.01, figure 3*a*), but there was no significant effect of exposure duration (*χ*^2^ = 0.280, df = 1, p > 0.05) and their interactions (*χ*^2^ = 3.491, df = 1, p = 0.062). Specifically, 38C3d pupae were four times more likely to develop deformed wings compared to the controls (z = −3.367, p < 0.01), while 38C5d pupae were nearly seven times more likely of developing deformed wings (z = −3.920, p < 0.001). Pupae of the 36C7d treatment showed a marginally significant, twofold increase in deformed wings (z = −2.482, p = 0.052). Pairwise comparisons between the other treatment groups and controls were not significant (36C5d: z = 0.656, p > 0.05; 37C3d: z = −2.001, p > 0.05; and 37C5d: z = −2.279, p > 0.05).

**Figure 3.**
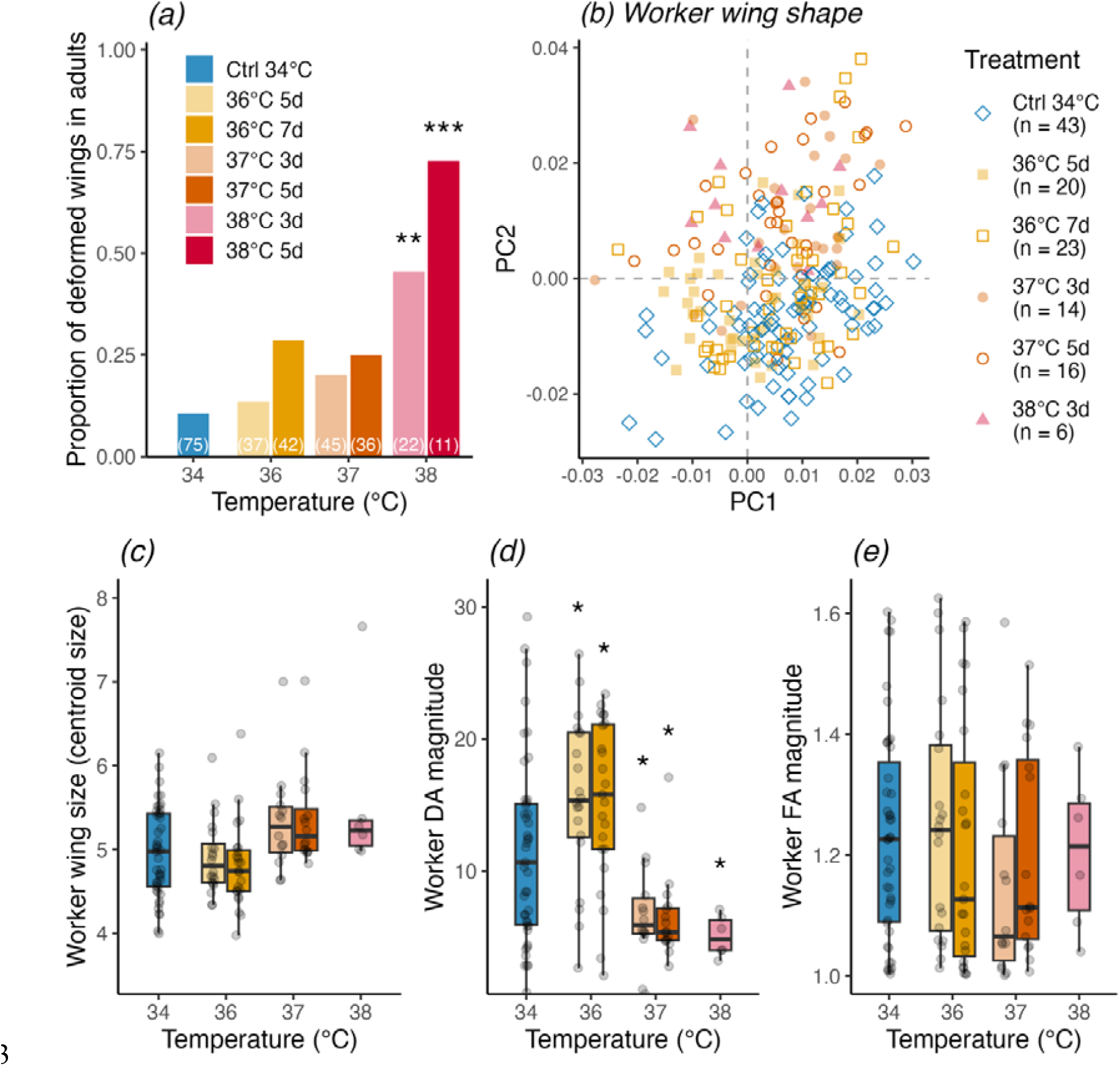
Effects of thermal stress on wing development in *B. terrestris*. *(a)* Proportion of deformed wings in emerged adult bees (including both males and workers) for each treatment group (sample sizes are indicated by white numbers). Deformed and male wings presented in *(a)* were excluded for detailed wing geometric morphometric analyses *(b-e)*, therefore resulting in lower sample sizes. *(b)* Principal component analysis (PCA) of wing morphology in emerged workers with fully developed wings across thermal stress treatments (see legend, n = samples size). *(c)* Influence of thermal stress treatments on wing size (centroid size) and the magnitude of *(d)* directional and *(e)* fluctuating asymmetry in fully developed worker wings. Controls (34°C, blue) were compared to individuals exposed to increased temperatures: of 36°C for 5 days (light warm yellow) and 7 days (warm yellow), 37°C for 3 days (light reddish-orange) and 5 days (reddish orange), and 38°C for 3 days (light red). Asterisks denote significant differences (* p < 0.05, ** p < 0.01, *** p < 0.001).

Our geometric morphometrics wing analysis, excluding deformed wings, indicated a significant effect on worker wing shape (PERMANOVA: F = 11.677, R^2^ = 0.197, df = 5, p < 0.001, figure 3*b*). Pairwise comparisons revealed significant differences between all treatment groups and the controls (36C5d, F = 7.259, p < 0.05; 36C7d, F = 5.005, p < 0.05; 37C3d, F = 21.125, p < 0.05; 37C5d, F = 29.494, p < 0.05; 38C3d, F = 25.168, p < 0.05). Similarly, we found that exposure affected centroid wing size in workers (LM: F = 4.205, df = 5, p < 0.01, figure S3*c*). Pairwise comparisons indicated a marginally significant increased worker wing size of the 37C5d (t = −2.568, p = 0.058) and 38C3d (t = −2.394, p = 0.073) treatment groups compared to the controls.

Furthermore, our thermal stress treatment significantly affected the DA magnitude in worker wings (LM: F = 9.466, df = 5, p < 0.0001, figure 3*d*), but not FA magnitude (Kruskal-Wallis: *χ*^2^ = 3.903, df = 5, p > 0.05, figure 3*e*). Workers exposed to 36°C showed higher DA (36C5d, t = −2.520, p < 0.05; 36C7d, t = −2.536, p < 0.05), while workers exposed to 37°C (37C3d, t = 2.655, p < 0.05; 37C5d, t = 2.932, p < 0.05) and 38°C (38C3d, t = 2.500, p < 0.05) showed lower DA compared to controls.

## 3. Discussion

Our study provides compelling evidence that thermal stress temperature and duration significantly impact pupal development in *B. terrestris* with notable effects on emergence rate (figure 1*a*), developmental duration (figure 1*b*), lipid stores (figure 2*d*), and wing development (figure 3). These findings highlight the extreme vulnerability of the pupal stage to thermal stress, particularly at 38°C, which appears to be a critical threshold for development disruption.

Although our treatments are extreme, they reflect potential heatwave scenarios in Europe with temperatures exceeding 38°C for several days as reported in parts of the British Isles, Mediterranean and Eastern Europe [38-40]. We are aware that our *in vitro* experiment oversimplifies the thermal stress that bumblebee pupae would experience naturally, especially in ground-nesting species such as *B. terrestris*, where ambient temperatures are buffered [20]. However, bumblebee species that typically nest above ground (e.g. *B. hypnorum* and *B. pascuorum*) or commercially managed colonies (e.g. *B. terrestris*), widely used for pollination services in greenhouses [21], are potentially more directly exposed to extreme thermal stress over prolonged periods. Furthermore, we expect more cold-adapted bumblebee species such as *B. lapidarius, B. alpinus*, and *B. poralris* to be even more vulnerable to thermal stress than *B. terrestris*, given its relative high heat tolerance [15, 19]. While natural colony responses to thermal stress, such as fanning, aim to cool nest temperatures, these thermoregulation efforts may negatively affect colony fitness and could be overwhelmed or fail under prolonged periods of stress [34, 35]. Here, we used *B. terrestris* as a model species to gain a mechanistic inside into the dose-response to thermal stress on pupal development by isolating them from colony.

Most notably, our findings show that emergence rates decreased by almost twofold and fivefold in pupae exposed to 38°C for 3 and 5 days, respectively, compared to controls (figure 1*a*). Despite *in vitro* rearing of L4 larvae and pupae deviates from natural nest conditions, the high emergence rate of controls suggests that our experimental setup allowed for normal development. While 38°C exposure slightly prolonged pupal development, a 36°C exposure slightly accelerated it (figure 1*b*), suggesting a potential developmental benefit in pupae to temperature elevation up to 37°C, followed by severe disruption at 38°C.

We observed no clear effects on worker body size proxies (figures 2*a-c*), except for an increased worker head width at 38°C. These results contrast with previous studies showing that colonies exposed to prolonged thermal stress would produce smaller offspring, indicated by smaller ITD [30, 32] and body mass [33], potentially reflecting resource allocation strategies such as reduced larval feeding. Alternatively, thermally stressed L4 larvae may have a decreased body weight gain due to reduced food consumption [28]. Despite potential limitations in statistical power due to the relative low emergence rates and resulting small sample sizes (figure 1*a*), particularly for the 38°C treatments, we speculate that larger pupae have a selective advantage under thermal stress (figure 1*b*, figure S3*a*). Although small insect body sizes should generally be favoured under warmer environmental conditions over the long term according to Bergmann’s rule [8], this rule does not apply to bumblebees [31].

In addition, we found that individuals exposed to thermal stress during pupal development (37°C for 5 days, 38°C for 3 and 5 days) had increased levels of the relative lipid content (fat body) as adult workers (figure 2*d*). This may indicate a potentially higher thermal stress resilience due to increased energy reserves, immunocompetence, and heat shock protein (HSP) synthesis [27, 50-53]. In fact, the insect fat body is the primary site for HSP synthesis in response to thermal challenges by activating stress-response pathways that upregulate the production of these chaperones to mitigate protein damage [50, 53]. Interestingly, the relative lipid content was not affected in workers that were exposed to thermal stress as L4 larvae, further underscoring the differential vulnerability of developmental stages [28].

Wing development was severely impaired by thermal stress temperature, notably with a fourfold and sevenfold increase in wing deformities in emerged adult bees from pupae exposed to 38°C for 3 and 5 days, respectively (figure 3*a*). Even excluding individuals with these deformities, workers wing shape altered for all treatments (figure 3*b*), which appears to be a natural response in colonies under thermal and parasitic stress [31, 54]. However, our data shows only marginal effects on worker wing size (figure 3*c*) with a similar pattern as for the other worker body size proxies (figure 2*a-c*). DA was affected by temperature in workers, with increasing DA at 36°C and decreasing DA at 37°C and 38°C (figure 3*d*), potentially also reflecting the different selection pressures among treatments. While FA remained unaffected in workers (figure 3*e*), confirming previous studies in honeybees [55] and bumblebees [31].

In conclusion, our *in vitro* study provides novel insights into the direct impact of varying thermal stress scenarios on *B. terrestris* pupal development. While our experimental design sacrifices ecological realism, it enabled precise control over timing and temperature. Our findings highlight a critical thermal stress threshold at 38°C, where severe disruptions in bumblebee pupal development manifest by particularly impairing emergence and wing development. Given the heat tolerance of *B. terrestris*, we expect more cold-adapted and above ground nesting bumblebee species are potentially more vulnerable to increasingly more frequent and severe heatwaves as climate change continues.

## Acknowledgments

[Anonymous]

## Author contributions

[Anonymous]

## Data accessibility

Datasets and R script used for statistical analysis and producing all figures have archived in Dryad repository: http://datadryad.org/share/vpCETsQuLDiGFIHSwioKRfRjwKhWotnYFRwZ72ovPC8

## Ethics statement

This study was conducted in accordance with the ethical regulations of the German Animal Welfare Act (TierSchG) for conducting experiments with insects.

## Funding

This study was carried out without any third-party funding.

## Competing interests

The authors have no competing interests.

## Notes

### Competing Interest Statement

The authors have declared no competing interest.

### Summary of Updates

We made a major revision on our original manuscript to improve the framing and clarity of our study.

