## Supplementary Information for "Impact of thermal stress during pupal development in a key pollinator"

[Anonymous]

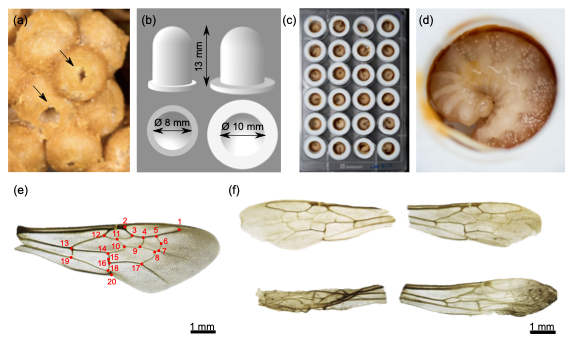

**Figure S1. *In vitro* larval rearing of L4 (a-d) and wing morphology analysis (e, f) in *B. terrestris*.** (a) L4 larvae were identified by the small opening for food provisioning (black arrows) and carefully collected. (b) Design of 3D-printed polylactide (PLA) artificial brood cells (capacity: 0.6 ml or 0.96 ml). (c) L4 larvae inside artificial brood cells in a 24-well plate. (d) Observation of L4 feeding on medium (see also supplementary video 1). (e) Fully developed right forewing showing the positions of the 20 landmarks (red dots) used for the geometric morphometric analysis, including centroid size, wing shape, directional asymmetry (DA), and fluctuating asymmetry (FA). (f) Representative examples of left and right forewings from two emerged adults with slight (top) and more severe (bottom) wing deformities. Individuals with deformed wings were excluded from further geometric morphometric analyses. Scale bar = 1mm.

**Table S2.** Overview of model structure and selection procedure based on lowest Akaike information criterion (AIC) for models meeting the model assumption criteria (see Supplementary RMD output for complete selection procedure and analysis). The best model fit is highlighted in bold. If criteria were not met, treatment effects were analysed using a non-parametric Kruskal-Wallis rank sum test (not listed here).

| **Response ^a^** | **Candidate models ^b^** | **Error structure** | **df** | **AIC** |
| --- | --- | --- | --- | --- |
| **emerged** | **~ temp*dur + (1\|colony) + (1\|collected)** | **binomial(link = "logit")** | **6** | **750.0324** |
|  | ~ temp + dur + (1\|colony) + (1\|collected) | binomial(link = "logit") | 5 | 776.4345 |
|  | ~ temp + (1\|colony) + (1\|collected) | binomial(link = "logit") | 4 | 775.9522 |
|  | ~ dur + (1\|colony) + (1\|collected) | binomial(link = "logit") | 4 | 809.5120 |
| **rel_lipid_content** | ~ temp * dur + sex + (1\|colony) + (1\|collected) | beta_family(link = "logit") | 8 | -889.7464 |
|  | ~ temp * dur + (1\|colony) + (1\|collected) | beta_family(link = "logit") | 7 | -887.7491 |
|  | **~ temp + dur + sex + (1\|colony) + (1\|collected)** | **beta_family(link = "logit")** | **7** | **-890.4072** |
|  | ~ temp + dur + (1\|colony) + (1\|collected) | beta_family(link = "logit") | 6 | -888.6448 |
|  | ~ temp + (1\|colony) + (1\|collected) | beta_family(link = "logit") | 5 | -889.5505 |
|  | ~ dur + (1\|colony) + (1\|collected) | beta_family(link = "logit") | 5 | -883.7733 |
|  | ~ sex + (1\|colony) + (1\|collected) | beta_family(link = "logit") | 5 | -886.5302 |
| **wings_deformed** | ~ temp*dur + sex + (1\|colony) + (1\|collected) | binomial(link = "logit") | 7 | 276.3331 |
|  | **~ temp*dur + (1\|colony) + (1\|collected)** | **binomial(link = "logit")** | **6** | **276.3267** |
|  | ~ temp + dur + (1\|colony) + (1\|collected) | binomial(link = "logit") | 5 | NA |
|  | ~ temp + (1\|colony) + (1\|collected) | binomial(link = "logit") | 4 | 276.3267 |
|  | ~ dur + (1\|colony) + (1\|collected) | binomial(link = "logit") | 4 | 285.2438 |

^a^ response variables: *emerged* =bees reaching adulthood; *rel_lipid_content* = lipid content in adults relative to their dry mass; *wings_deformed* = bees with deformed wings

*^b^ candidate models: temp* = treatment temperature; *dur =* exposure duration; *colony* = colony ID; *collected* = collection cohort

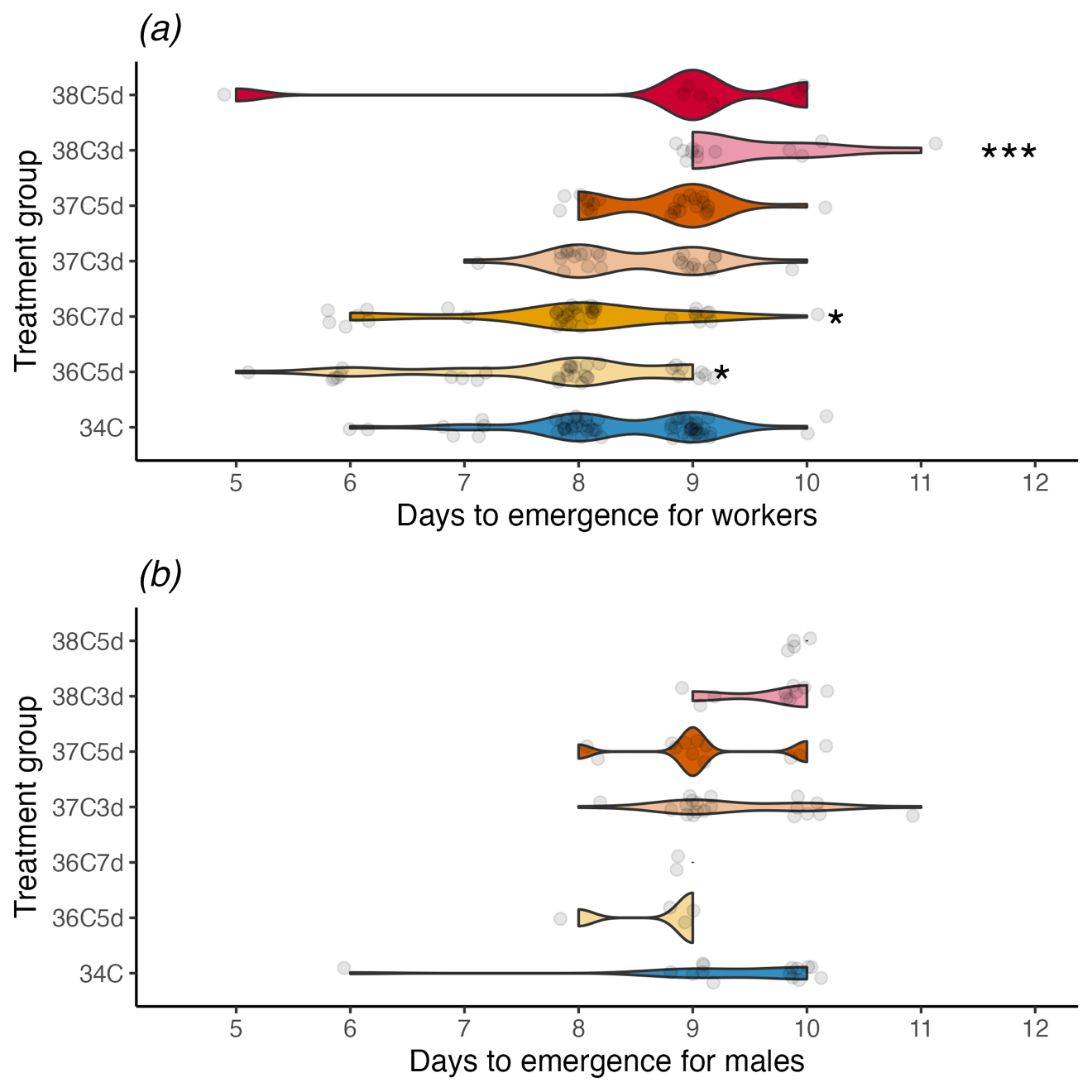

**Figure S2. Direct effects of varying thermal stress temperatures and durations on (a) worker and (b) male developmental duration in *B. terrestris* pupae.** Controls (Ctrl, reared at constant 34°C, blue) were compared to individuals exposed to increased temperatures: of 36°C for 5 days (light warm yellow) and 7 days (warm yellow), 37°C for 3 days (light reddish-orange) and 5 days (reddish orange), and 38°C for 3 days (light red) and 5 days (deep red). Significant differences are marked by asterisks (* p < 0.05, ** p < 0.01, *** p < 0.001).

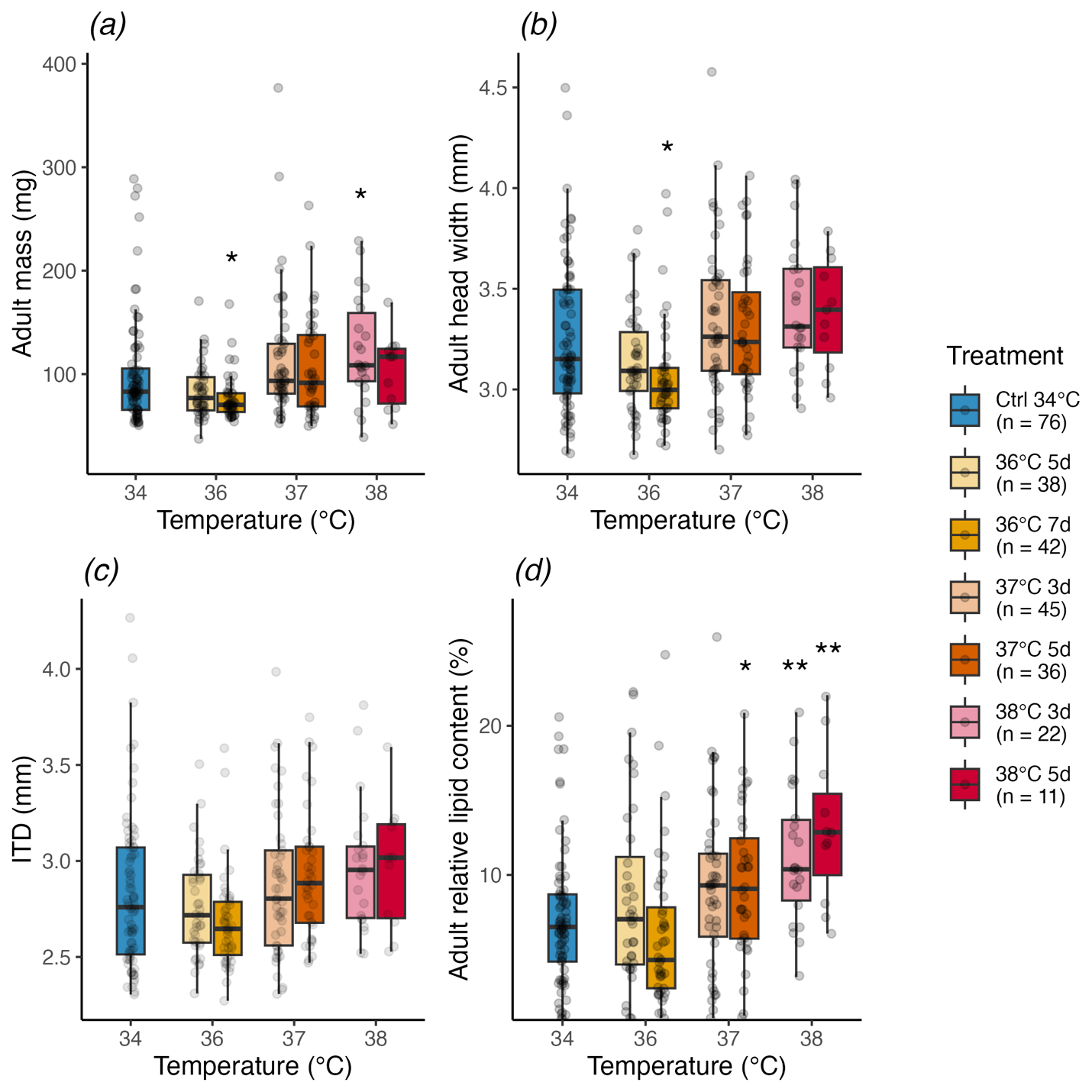

**Figure S3. Effects of thermal stress temperature and duration during *B. terrestris* pupal development on overall adult morphometrics and lipid content (including both male and worker data):** *(a)* adult mass, *(b)* head width, *(c)* intertegular distance (ITD), and *(d)* relative lipid content in 2-day-old adults. Controls (34°C, blue) were compared to individuals exposed to increased temperatures: of 36°C for 5 days (light warm yellow) and 7 days (warm yellow), 37°C for 3 days (light reddish-orange) and 5 days (reddish orange), and 38°C for 3 days (light red) and 5 days (deep red). Sample sizes (n) are provided in the legend. Asterisks denote significant differences (* p < 0.05, ** p < 0.01).
